# Cell surface remodeling caused by the loss of flippase subunit TMEM30A in immune cells

**DOI:** 10.64898/2026.02.21.707187

**Authors:** Cenk O. Gurdap, Franziska Ragaller, Marion Muller, Ellen Sjule, Taras Sych, Linnea Blomén, Fredrik Thoren, Ilya Levental, Kandice R. Levental, Quentin Sattentau, Erdinc Sezgin

## Abstract

Plasma membrane lipid asymmetry is tightly regulated and fundamental to mammalian cell physiology. TMEM30A is the β-subunit of P4-ATPases, flippase enzymes that maintain strict phosphatidylserine (PS) asymmetry by pumping it from the outer to the cytosolic leaflet. Loss of TMEM30A function causes constitutive PS externalization and has been implicated in diseases such as diffuse large B-cell lymphoma and tumour immune evasion. Here, we systematically define the biophysical and molecular consequences of TMEM30A deletion in immune cells. Using a live-cell lipid reporter, membrane order probe, and surface proteome mapping, we show that TMEM30A-knockout cells display robust PS externalization accompanied by faster lateral diffusion of membrane constituents and decreased plasma membrane order. Surface proteome reorganization includes increased abundance of tetraspanins and CD47. Further, TMEM30A loss triggers glycocalyx remodeling via ADAM10-dependent shedding that removes major transmembrane mucins, including CD43 and CD162. Together, these data reveal a coordinated reorganization of lipids, glycans, and proteins upon TMEM30A loss, suggesting mechanistic links between flippase dysfunction and increased plasma membrane dynamics and potential sensitization to immune therapy. Furthermore, our study provides an integrated surfaceome framework that might shed light on the relationship between TMEM30A expression and clinical outcomes in cancer.

## Introduction

Plasma membrane asymmetry is highly regulated and essential for numerous cellular processes. Phosphatidylserine (PS) and phosphatidylethanolamine (PE) are typically confined to the inner leaflet, while phosphatidylcholine and sphingomyelin are more abundant on the outer leaflet^1,2^. This asymmetric organization is crucial for, amongst other functions, maintaining membrane curvature, enabling vesicle formation, and facilitating signal transduction^2^. Disruption of this asymmetry, such as via the externalization of PS, can serve as a signal for immune recognition of apoptotic cells^3^. Additionally, homeostatic lipid distribution supports the function of membrane proteins and receptors, ensuring effective interaction and transport across the membrane^3^. Thus, membrane asymmetry is fundamental to cell viability and cell-cell interactions^4^. However, under certain physiological and pathological conditions, such as during apoptosis, cell and platelet activation, or cellular stress, PS is externalized to the outer leaflet^3^. In apoptosis, PS exposure acts as an “eat-me” signal^5^, recognized by phagocytes to facilitate the non-inflammatory clearance of dying cells^6^. Moreover, externalized PS is associated with activation of “A Disintegrin And Metalloprotease” (ADAM)-family metalloproteases^7,8^, which can cleave multiple substrates. Amongst these ADAM substrates are glycocalyx-embedded transmembrane mucins. Their shedding from the surface of apoptotic immune cells increases recognition of the cells by phagocytes^9^. Additionally, PS can also play immunomodulatory roles under other, non-apoptotic conditions^10^. These roles suggest a complex interplay between PS and other surface molecules.

The strict maintenance of PS on the inner plasma membrane leaflet is mediated by several enzymes. TMEM30A, also known as CDC50A, is an essential subunit of the P4-ATPase phospholipid flippases, which maintain the asymmetric distribution of phospholipids across cellular membranes^11,12^. Cells lacking functional TMEM30A show constitutive PS flopping to the outer leaflet and are better at evading natural killer (NK) cell-mediated cytotoxicity^13^. Relatedly, recurrent biallelic loss-of-function mutations in TMEM30A were identified in diffuse large B-cell lymphoma (DLBCL), an aggressive cancer, through genomic and transcriptomic analysis of patient samples from a population-based registry^14^. TMEM30A was suggested as a putative tumor suppressor, potentially changing membrane organization in B cells. However, the role of TMEM30A function in pathology is complicated, since TMEM30A loss has also been correlated with favorable clinical outcomes in cancer treatment^14^. TMEM30A-deficient cells had higher uptake of chemotherapy drugs, and TMEM30A-deficient tumors exhibited increased infiltration of tumor-associated macrophages and showed enhanced sensitivity to anti-CD47 therapy^14^. While these results suggest that loss of TMEM30A changes the surface properties of cells, the biophysical and molecular changes on the cell surface, and how they relate to therapy outcome, are not clear.

Here, we investigated the protein, carbohydrate, and lipid remodeling taking place at the plasma membrane of TMEM30A knock-out (TMEM30A-KO) immune cell lines. We show that TMEM30A-KO immune cells exhibit elevated PS on the outer leaflet, faster diffusion of molecules in the plasma membrane, and reduced plasma membrane order. Surface protein mapping experiments showed that KO cells differentially regulate several proteins including upregulation of members of the membrane protein-organizing family of tetraspanins, and the checkpoint molecule CD47, and downregulate transmembrane mucins CD43 and CD162. Using ADAM10 inhibitors, we show that increased PS exposure correlates with ADAM10-dependent cleavage and loss of mucins, in particular CD43, from the surface. Overall, our data reveal a major reorganization of lipids, carbohydrates, and proteins on the tumor immune cell surface upon loss of TMEM30A, suggesting potential explanations for the physiological phenotypes of TMEM30A mutations, such as immune evasion, enhanced membrane mobility, increased surface accessibility, and better responses to anti-CD47 therapy.

## Results and Discussion

To address how the loss of TMEM30A changes the cell surface with implications for intercellular interactions and immune evasion, we initially characterized the plasma membrane biophysical properties of two cell types in which TMEM30A was knocked out.

### TMEM30A-KO cells expose PS on the cell surface

PS externalization has previously been suggested as a protection mechanism against immune cell killing^15^. To this end, we first studied the PS exposure on two immortalized immune cell lines: the Jurkat T cell line derived from an Acute Lymphoblastic Leukemia patient and the K562 line derived from a Chronic Myeloid Leukemia patient. Wild-Type (WT) cells were compared with TMEM30A-KO for cell surface PS expression using fluorescently tagged Lactadherin C2 domains (LactC2)^16,17^ followed by confocal microscopy imaging and flow cytometric analysis. Both microscopy **(Fig. 1A)** and flow cytometry **(Fig. 1B, see Supporting Figure 1 for non-normalized replicates)** analysis showed that KO cells showed significantly higher LactC2 binding compared to WT, confirming that KO cells exhibit increased PS on the exoplasmic leaflet^13^.

**Figure 1.**
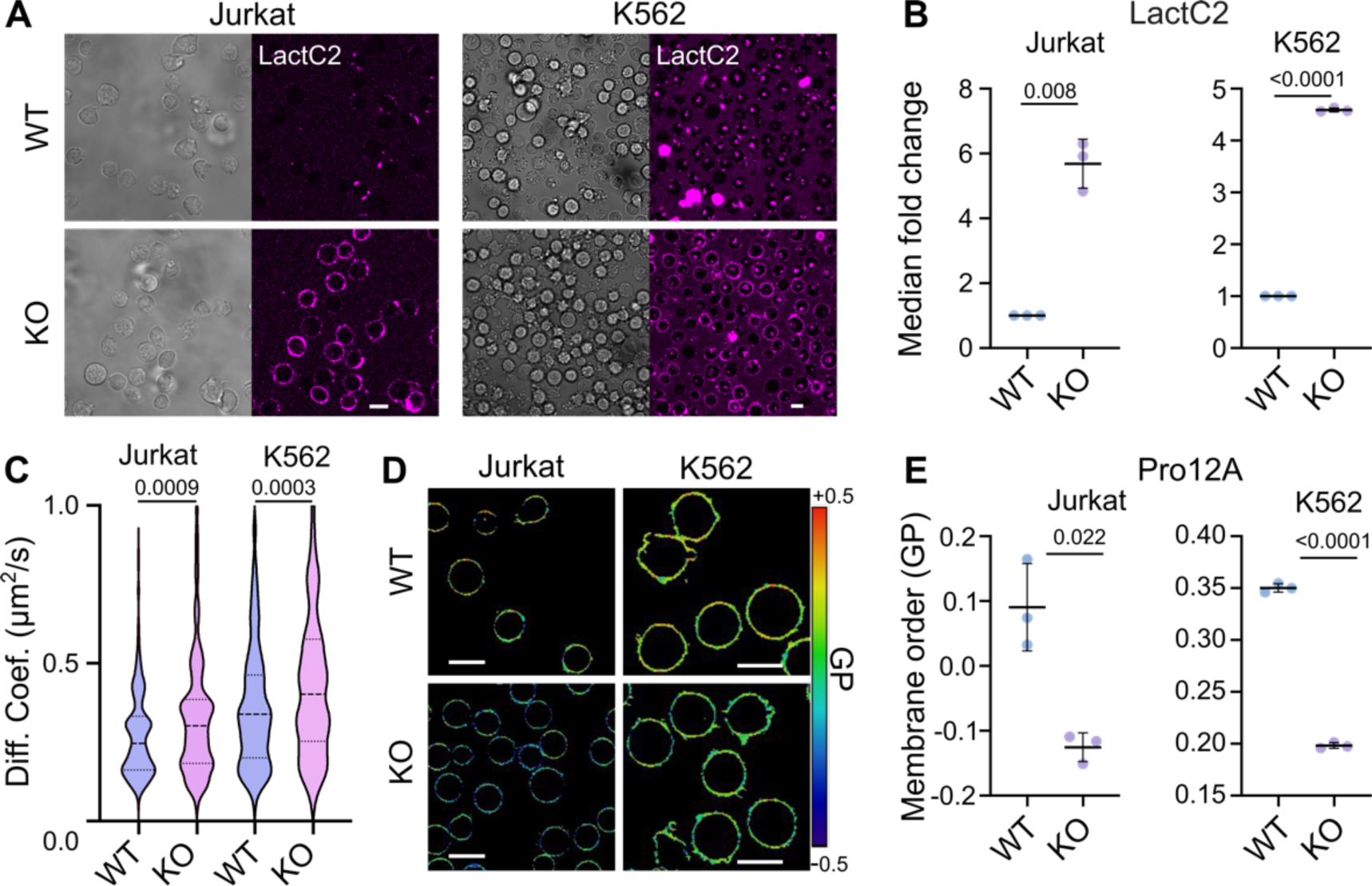
Biophysical properties of TMEM30A-KO cells. A) TMEM30A-KO Jurkat and K562 cells are viable and exhibit PS+ puncta on the cell surface. PS is labelled with LactC2-Alexa647. B) PS exposure was quantified in both cell types using flow cytometry, mean ± SD. The median fluorescence was derived for each sample using identical laser and detector settings for WT and KO lines. The median fluorescence intensities for WT replicates were normalized to 1 and the KO signal was represented as fold-change compared to WT. C) Diffusion of Abberior Star Red-PEG-DPPE is faster in TMEM30A-KO cells compared to WT cells. Violin plots showing the median and first and third quartiles. D) TMEM30A-KO cells have more fluid plasma membranes compared to WT cells measured by Pro12A shown as GP map. E) Membrane order (GP index) was quantified using spectral biophysical cytometry, mean ± SD. Each data point is an independent experiment (n=3) and p-values are calculated with a Mann-Whitney U test. Scale bars are 10 μm.

### Loss of TMEM30A causes biophysical remodeling of the plasma membrane

The composition and leaflet organization of lipids imparts functional membrane biophysical properties. For example, high membrane order has been proposed as a protection mechanism against immune cell killing^18^. Interestingly, loss of membrane asymmetry through PS scrambling has been shown to reduce membrane ordering^19^. Therefore, we tested whether loss of TMEM30A affects membrane order in the plasma membrane. We first investigated membrane fluidity by measuring the diffusion of an outer leaflet lipid tracer using fluorescence correlation spectroscopy (FCS). FCS measures the diffusion of molecules by detecting the fluorescence fluctuations due to molecular movement through the microscope observation volume. From the FCS curves, we measured the diffusion coefficient, where higher values mean faster diffusion. The diffusion of Abberior Star Red PEG-labelled 1,2-dipalmitoyl-sn-glycero-3-PE (DPPE) lipid tracer was significantly faster in KO cells compared to WT cells for both cell lines **(Fig. 1C)**, suggesting that loss of TMEM30A increases plasma membrane fluidity upon TMEM30A loss. This inference was further supported by membrane order measurements by spectral imaging^20^ and flow cytometry^21,22^ using the environment sensitive probe Pro12A^23^ in **(Fig. 1D-1E, see Supporting Figure 1 for non-normalized replicates)**. Pro12A reports membrane order by shifting its emission spectrum depending on the lipid environment, which can be ratiometrically quantified as Generalized Polarization (GP) value, where higher GP represents higher membrane order^24^. KO cells showed significantly less ordered membranes compared to WT cells in both Jurkat and K562 (colder colors in **Fig. 1D** and lower values in **Fig. 1E**). Of note, Pro12A specifically measures membrane order in the outer leaflet^23^, hence these results confirm that the outer leaflet of KO cells is more fluid than that of WT cells.

Regulation of the properties of the two leaflets as to whether they are independent from each other, is largely unexplored. Therefore, we also set out to measure the properties of the inner leaflet. To do this, we used Halo-Tag protein reporters in combination with NileRed-Halo^25^. These proteins localize the HaloTag to the inner leaflet of the plasma membrane, allowing NileRed-Halo to specifically access this membrane leaflet. The lifetime of NileRed is sensitive to the lipid environment, hence we could measure inner leaflet lipid order using this approach. Interestingly, unlike the outer leaflet, we did not observe significant differences between the inner leaflet membrane order of TMEM30A KO cells and WT cells **(Supporting Figure 2)**. The fact that TMEM30A-deficient cells maintain normal inner-leaflet lipid order while exhibiting decreased outer-leaflet order potentially suggests an asymmetry in how the plasma membrane responds to loss of phospholipid flippase activity and how two leaflets are regulated in cells. The stability of the inner leaflet implies that its biophysical properties are differently regulated compared to the outer leaflet. By preserving inner-leaflet order, TMEM30A KO cells may maintain functional integrity of pathways that depend on precise lipid packing and charge distribution, allowing them to continue proliferating despite substantial outer-leaflet perturbations. In contrast, mechanisms disrupting both leaflets can compromise inner-leaflet organization causing detrimental outcomes, including cell death. These findings support the idea that not all forms of phosphatidylserine externalization are biologically equivalent: TMEM30A loss selectively perturbs the outer leaflet while sparing the inner leaflet, thereby preserving cellular homeostasis even under conditions of chronic PS exposure.

Overall, these data show that the plasma membrane, specifically the outer leaflet, is substantially reorganized in KO cells, explaining the previously reported increased diffusivity of receptors on the cell surface^14^. However, our observation does not align with the immune evasion hypothesis^15^ since the disordering of the membrane in KO cells would be expected to sensitize, rather than protect, them from immune cell killing. To explain this effect and further characterize the cell surface changes consequent to loss of TMEM30A, we investigated the remodeling of specific plasma membrane proteins implicated in immune regulation.

### Loss of TMEM30A leads to major surface proteome remodeling

The surface proteome (e.g., type, abundance and organization of the plasma membrane proteins) is a major determinant of cell-cell interactions. Therefore, we tested whether the loss of TMEM30A can remodel the surface proteome, with potential implications for therapeutic responses to, and immune evasion by, cancer. Changes in membrane lipid composition and the loss of lipid asymmetry can profoundly influence the repertoire and function of surface proteins, since many of these rely on specific lipid interactions for proper localization, conformation, and/or activity^26^. For instance, PS is suggested to interact with and regulate the activity of ADAM family sheddases^8,9,27,28^ that cleave the extracellular domains of numerous membrane proteins. Disruption of lipid asymmetry and externalization of PS could therefore, directly or indirectly, lead to ADAM activation and thereby reshape the surface proteome. Moreover, loss of asymmetry can disrupt endocytic and exocytic trafficking routes^29^ potentially trapping certain proteins in endosomes or incorrectly recycling them back to the surface. Finally, changing PS leaflet distribution can affect the spatial arrangement of proteins including integrins, growth-factor receptors, and ion channels^30,31^. Thus, we set out to investigate how the absence of TMEM30A affects the cell surface protein landscape.

We measured the remodeling of the surface proteome using the ‘molecular pixelation’ assay (**Fig. 2A**)^32^. This assay works by labeling cell-surface proteins with antibody-oligonucleotide conjugates (AOCs) and then converting their physical proximity into DNA information that can be decoded by sequencing. After cells are fixed, DNA pixels (single-stranded DNA molecules) are added, which hybridize with the AOCs and associate them into local neighborhoods. By rolling circle amplification, the neighborhood information is added to the AOCs and the oligos are subsequentially amplified by PCR. This creates DNA sequences containing barcodes to identify the unique AOCs, protein identity and neighborhood membership. These DNA molecules are sequenced and computationally reconstructed to quantify protein abundance and colocalization and to create interaction networks of single cells with a spatial resolution of 280 nm, using standard next generation sequencing methods.

**Figure 2.**
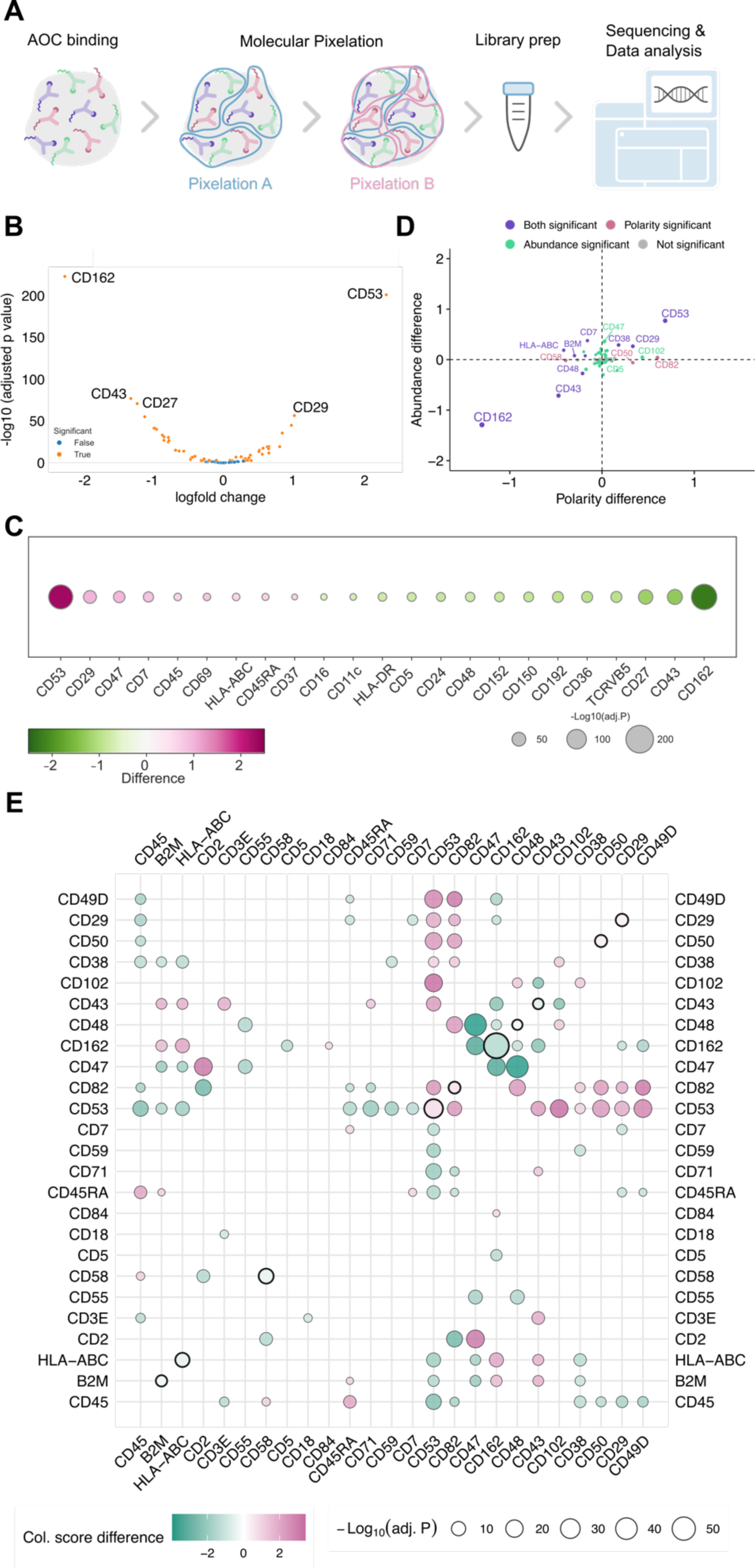
Jurkat surface proteome is dramatically reorganized by TMEM30A-KO cells. **A)** Scheme of molecular pixelation experiment. **B)** Volcano plot for the abundance of proteins differentially expressed on cell surface of WT vs. KO Jurkat cells. Orange dots indicate significantly up- or downregulated proteins (adjusted p-value < 0.01). **C)** Proteins of differential abundance (log-fold change >0.6) in WT vs. KO Jurkat cells. Color corresponds to log-fold change and size of the bubbles indicates the adjusted p-value. **D)** Proteins that are significantly regulated in polarity versus abundance. **E)** Differential colocalization map of relevant proteins (difference in colocalization in KO compared to WT).

Using antibodies specific for the selected proteins, we ran molecular pixelation analysis in Jurkat cells and selected the proteins whose surface abundance was most affected by TMEM30A-KO (**Fig. 2B, C, and Supporting Figure 3)**. Transmembrane mucins such as CD162 and CD43 were reduced significantly, while tetraspanin CD53 and integrin CD29 were significantly increased. Interestingly, CD47, a negative regulator of phagocytosis and immune killing, was also increased in KO cells. Besides abundance, polarity of the proteins (e.g., distribution of the proteins towards one side of the cell surface), a critical parameter for immune cell function^33^, also changed in KO cells. The tetraspanin CD53 was particularly highly polarized in KO cells. Reduced polarization is often correlated with reduced abundance as expected. When abundance and polarity were analyzed together **(Fig. 2D)**, mucins showed clear downregulation while tetraspanin CD53 and integrin CD29 showed obvious upregulation in KO cells. CD47 was significantly different in abundance but not in polarization, whereas the tetraspanin CD82 exhibited significantly different polarization but not abundance. Finally, we compared protein colocalization differences between WT and KO Jurkat cells (**Fig. 2E**). Purple color indicates increased colocalization of protein pairs in KO cells compared to WT cells. Tetraspanins CD53 increases in colocalization with CD82, mucins like CD43 and adhesion-associated molecules such as CD102, CD49D, CD29 and CD50. CD47 colocalization with CD2 also increased in KO cells. In contrast, CD162 interacts less with CD47 and other mucins including CD43. These data reveal that there is extensive surface proteome remodeling upon loss of TMEM30A. Some proteins increase either in abundance (e.g., CD47) or clustering (e.g. CD82), or increase in both abundance and clustering (e.g., CD53, CD29) while some decrease in abundance and clustering (e.g., CD162, CD43).

Increases in CD47 expression, which relays an inhibitory signal upon interaction with immune cells such as macrophages and NK cells^34^, might potentially contribute to the immune cell evasion of cells with non-functional TMEM30. Moreover, this increase might account for the susceptibility of TMEM30A-deficient DLBCL tumors to anti-CD47 therapies. Changes in tetraspanin expression regulate interactions between surface proteins, specifically with ADAMs modulating their enzyme activity and specificity^35–37^. For example, tetraspanin CD53 interacts with ADAM metalloproteases^38^ and therefore variation in the expression of tetraspanins such as CD53 and CD82 should be further investigated in the context of ADAM activity. In this context, mucins were found to be especially differentially regulated. Interestingly, downregulation of overall glycosylation was also observed in adherent cells with flippase knock-down adherent cells^39^. Therefore, next we performed a more mechanistic analysis on mucin alterations in TMEM30A-KO cells.

### Surface mucins are cleaved by ADAM10 in TMEM30A-KO cells

Mucins are highly glycosylated cell surface glycoproteins that contribute heavily to the glycocalyx, the interactive barrier between cells and their environments^40^. This mucin barrier is crucial for protecting cells from infection and damage, but also selectively regulates interactions with other cells. Loss of mucins has been shown to affect intercellular interaction with pathogens^41^, immune cells and small molecules^42–44^. During apoptosis, the externalization of PS is associated with the activation of metalloprotease ADAM10 to cleave certain substrates from the surface, including mucins^8,9,27,28^.

Previous reports showed that TMEM30A-deficient cells are more accessible to small molecule drugs and to phagocyte recognition^14^ suggesting changes to the glycocalyx. Taking these observations together, we tested whether loss of TMEM30A causes ADAM10-dependent reorganization of the mucin barrier at the surface of immune cells. To this end, we used flow cytometry to test the cell surface abundance of specific mucins that were previously shown to be cleaved by ADAM10 in T-cells, namely CD43, CD162 and MUC1. As a control, we also studied two mucins that are not cleaved by ADAM10, namely CD45 and MUC24. Consistent with results from the molecular proximity assay, we observed significantly reduced levels of CD43 and CD162 and MUC1 in TMEM30A-KO cells, but no significant differences for CD45 and MUC24 **(Fig. 3A)**. To confirm that this decrease is due to ADAM10 activity rather than trafficking or transcriptional downregulation, we incubated Jurkat cells with the ADAM10-specific inhibitor (GI254023X, shortened to GI) or an ADAM10 and −17 inhibitor (GW 280264X, shortened to GW) at a non-toxic concentration (10 μM) for 24 h, and measured mucin loss. We observed a complete reversion of the loss of ADAM10-cleaved mucins after treatment of KO cells with the inhibitors, confirming the ADAM10-dependence of mucin shedding in TMEM30A-KO cells **(Fig. 3B)**. Of note, there was also a slight increase in the mucin signal in WT cells **(Supporting Figure 4)**, suggesting a basal level of ADAM10 activity and shedding of the extracellular domains of mucins even without externalized PS, which was suggested recently^28^, but may also correspond to constitutive low levels of PS expression at the outer leaflet of the WT plasma membrane (e.g., **Fig. 1A**).

**Figure 3.**
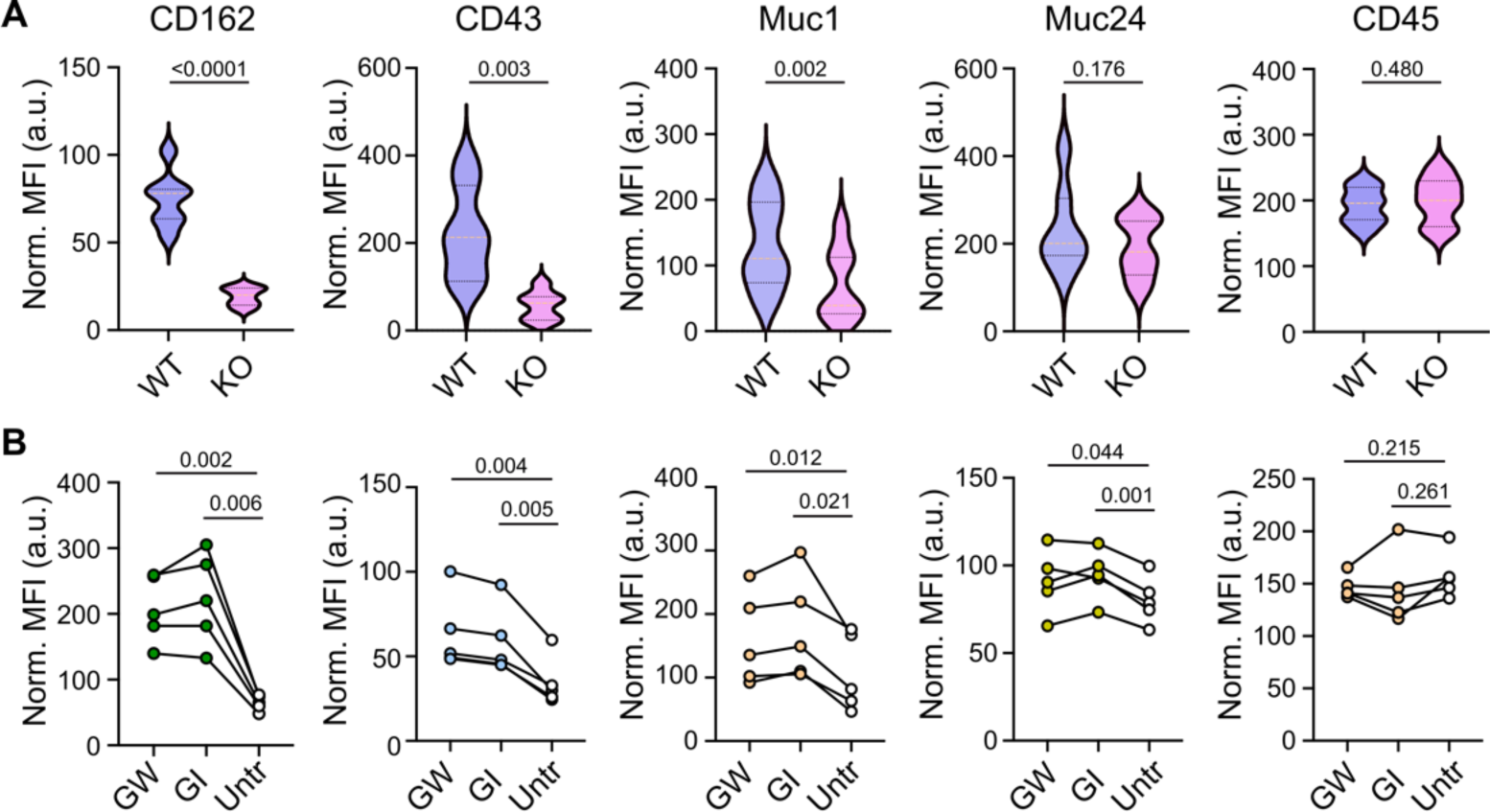
Mucin profiling in TMEM30A-KO Jurkat cells. **A)** Flow cytometry analysis of mucin abundance on cell surface of TMEM30A-KO Jurkat cells. Data expressed as violin plots of mean fluorescence intensity. **B)** Inhibition of mucin shedding shown by flow cytometry analysis of mucins in KO cells treated with ADAM10 inhibitor (GI) or an ADAM10/17 inhibitor (GW). Data points represent means of independent experiments; p-values calculated using RM one-way ANOVA test (n=5 independent experiments).

These data reveal mucin shedding from the surface of TMEM30A-KO cells in a PS- and ADAM10-dependent manner. Reduced mucin abundance implies disruption of the mucin barrier within the glycocalyx and increased accessibility of cells to external interactions and stimuli.

### Conclusion

TMEM30A-KO immune cells display robust PS externalization, as confirmed by LactC2-based flow microscopy and cytometry indicating loss of membrane asymmetry. This lipid redistribution triggers substantial biophysical remodeling of the plasma membrane: KO cells exhibit significantly increased membrane fluidity, demonstrated by faster diffusion of a lipid tracer in FCS measurements, and markedly decreased membrane order as quantified by Pro12A spectral imaging and spectral flow cytometry. To determine whether these changes extend to the protein landscape, we profiled the surface proteome using molecular pixelation and found extensive remodeling, including strong downregulation of CD162 and CD43 mucins and upregulation of tetraspanins (e.g. CD53), integrin CD29, and the immune checkpoint protein CD47. In addition to altered abundance, protein polarity and colocalization networks were reorganized, with increased clustering of CD53 and altered interactions involving CD47. Mechanistically, we demonstrated that mucin loss was driven predominantly by ADAM10-dependent shedding. Together, these findings reveal that TMEM30A loss induces a coordinated remodeling of surface lipids and proteins, altering membrane biophysics and driving loss of otherwise highly expressed membrane mucins.

Our results provide a potential mechanistic framework that help to explain why loss-of-function TMEM30A mutations - despite being tumor-promoting - are consistently associated with improved therapeutic responses, including enhanced sensitivity to chemotherapy and superior outcomes with anti-CD47 immunotherapy. We show that loss of TMEM30A leads to constitutive PS exposure on the outer membrane associated with ADAM10-dependent shedding of major surface mucins is anticipated to dilute the dense glycocalyx barrier that normally restricts drug penetration and shields tumor cells from immune engagement. Of particular importance, CD43 has very recently been described as the major leukemic glycocalyx molecule inhibiting macrophage phagocytosis and NK and T cell-mediated killing^45^. On one hand, reduced CD43 levels, as seen here, would limit this immune-resistance activity, in principle promoting tumor cell clearance. On the other hand, decreased glycocalyx density imposed by ADAM10-mediated mucin loss would more generally increase cell surface accessibility to soluble molecules, offering a possible explanation for the previously reported heightened uptake of chemotherapeutic agents in TMEM30A-deficient lymphomas. In parallel, we found that TMEM30A-KO cells upregulate CD47 on their surface, creating an expanded and more exposed “don’t-eat-me” checkpoint landscape. This molecular reorganization makes CD47 more available for antibody binding. Additionally, the resulting surfaceome remodeling including loss of the mucin barrier may render tumor cells more penetrable to drugs and more responsive to immunotherapy. Together our results highlight the importance of revealing the surface architecture of diseased cells to develop better therapies against them.

The increased B-cell receptor (BCR) mobility reported in TMEM30A-deficient tumors could also be explained by our biophysical data. By demonstrating that TMEM30A loss increases lateral diffusion and reduces membrane order of the outer leaflet, we provide a potential physical basis for the faster cell surface BCR movement that was reported previously^14^. In membranes, higher fluidity theoretically accelerates receptor encounters with coreceptors and kinases, lowers the energetic barrier for nano-cluster formation, and shortens the time to productive signaling assemblies^46^. Thus, the disordering of the lipid bilayer driven by PS externalization and consequent remodeling might create a membrane environment that is more permissive to BCR diffusion, clustering, and activation. In practical terms, this might imply that TMEM30A-deficient B cells might be intrinsically more prone to initiate and sustain signaling due to diffusion-controlled interactions in the plasma membrane, mechanistically aligning our measurements with the observed increase in BCR mobility and signaling propensity in TMEM30A-mutant tumors. However, such hypothetical mechanistic links need to be tested in the future.

Strong negative surface charge, high membrane order, a thick glycocalyx, and the presence of inhibitory ligands are classically associated with immune evasion. TMEM30A-deficient cells were previously reported to evade NK cell recognition by triggering inhibitory TIM-3 receptors on NK cells^13^. We extend these findings and show that TMEM30A-KO cells display a more complex phenotype and exhibit features consistent with both reduced and increased susceptibility to NK-cell–mediated killing. Reduced NK cell mediated killing may be promoted by pronounced PS-dependent surface negativity and a marked upregulation of CD47, which, at least in murine systems, strengthens the “don’t-kill-me” inhibitory signal. This, associated with membrane disordering should, in principle, increase accessibility to immune cells. These apparently opposing outcomes highlight that immune recognition is a multi-parametric process shaped by the integrated contributions of many surface cues.

Finally, our surfaceome mapping reveals extensive remodeling of protein abundance, clustering, and colocalization networks, suggesting that additional organizational features beyond individual markers critically shape immune cell-target cell interactions. Together, these findings suggest that clinically relevant TMEM30A phenotypes might arise from a complex interplay of lipid, protein, and glycocalyx changes, underscoring the need for future studies to dissect how these parameters collectively modulate immune synapse formation and cytotoxic responses.

Experimentally controlled PS externalization is not trivial to achieve. Our system leads to PS externalization due to the loss of flippase subunit TMEM30, which might have other unforeseen effects on cells. In related settings, PS exposure triggered by apoptosis was investigated; however, apoptosis also changes many other cellular processes besides PS exposure. Therefore, the findings here and in apoptosis-related studies should be tested using other PS externalization mechanisms in well-controlled model systems. For example, several groups have shown that treating cells with cyclodextrins^19^, ionophores^6^ or oxidized lipids^47^ leads to PS externalization and outer leaflet reorganization. Such systems, in combination with advanced imaging^48^, can be utilized in the future to elucidate the role of lipid asymmetry in cell-cell interactions including in cancer cell immune evasion and drug resistance.

## Materials and Methods

### Cells

Wild type and KO cells were described in detail in ref ^13^. Briefly, gene knockouts in Jurkat cells were generated using a ribonucleoprotein (RNP)-based CRISPR–Cas9 approach, whereas Cas9-expressing K562 cells were edited using both RNP- and plasmid-based methods. Guide RNAs targeting TMEM30A were selected from validated sources (Broad Institute or IDT) and verified using Synthego tools. For RNP editing, gene-specific crRNAs were annealed with Alt-R tracrRNA (IDT) and complexed with Alt-R S.p. HiFi Cas9 nuclease V3 to form RNPs. Cells were electroporated using the Neon transfection system (Invitrogen). In K562 cells, knockouts were generated in a Cas9-expressing background. Cells were sorted 48 h post-transfection based on fluorescent reporter expression or Annexin V staining (for TMEM30A), expanded, and validated by PCR and Sanger sequencing, with indel analysis performed using Synthego ICE.

For plasmid-based editing in K562 cells, guide RNAs were cloned into fluorescent reporter plasmids (mEGFP-C1 or EBFP2-C1), which were transfected following standard cloning, amplification, and purification procedures. All cells were mycoplasma negative.

### Confocal imaging, spectral imaging and spectral flow cytometry

*PS measurement with LactC2 labeling:* Cells seeded two days prior were harvested, counted, and distributed into Eppendorf tubes at a density of 300,000 cells per 1 mL RPMI medium. PS exposure was assessed using Bovine Lactadherin–Alexa Fluor 647 (PROLYTIX) and analyzed by flow cytometry. A total of 2 µL of 83 µg/mL dye was added to each sample and incubated for 15 minutes at room temperature (RT). Cells were then washed twice with PBS and finally resuspended in 200 µL PBS before transfer to flow cytometry tubes. Data acquisition was performed using 640 nm excitation (red laser) and emission detection at 670–688 nm (center 670 nm, 18 nm bandwidth; R2 channel). Analysis was conducted in FCS Express. The median autofluorescence signal from unlabelled cells was subtracted from labelled samples. Median fluorescence intensities were then used to calculate fold changes between WT and KO cells.

### Membrane order with Pro12A labeling

Spectral imaging of cells was performed as described in the protocol in ref^20^. Briefly, spectral confocal imaging was performed to assess membrane organization using the polarity-sensitive probe Pro12A. Cells were stained with Pro12A (final concentration 200 nM) for 2-5 min at room temperature, followed by washing in phenol red–free imaging medium. Imaging was conducted using a confocal laser scanning microscope equipped with a spectral detector and a 40X water objective. Pro12A was excited with a 405-nm laser line, and emission spectra were collected across the visible range using lambda mode. Spectral images were processed to extract fluorescence emission profiles on a pixel-by-pixel basis. Generalized polarization (GP) analyses were performed to quantify membrane order, using 440 and 500 nm as ratiometric wavelengths. Image analysis was performed using scripts as describes previously^20,49^.

Spectral flow cytometry was performed as described^22^. Briefly, 300,000 cells per sample were prepared as described above and washed twice with PBS. Cells were transferred to flow tube with a final volume of 300 uL PBS and labelled with 0.4 µL of 200 µM Pro12A dye for 1 minute at RT immediately prior to acquisition. Flow cytometric analysis was performed using 405 nm excitation (violet laser). Membrane order was quantified by calculating the generalized polarization (GP) value using V1 (420–435 nm, center 428 nm, 15 nm bandwidth) and V7 (533–550 nm, center 542 nm, 17 nm bandwidth) channels. Details of gating strategy, acquisition parameters, and data analysis are provided in ref^21,22^.

For each replicate for microscopy >10 images were analyzed, and each image contains >10 cells. For spectral flow cytometry, each replicate measured >5000-10000 cells.

### Fluorescence Correlation spectroscopy

Diffusion of Abberior Star Red-PEG-DPPE (Abberior) was measured using Fluorescence Correlation Spectroscopy (FCS). Cultured cells were collected, the culture medium was replaced with Leibovitz’s L-15 serum-free medium and supplemented with 50 nM of Abberior Star Red-PEG-DPPE. A Zeiss LSM 780 microscope with 40 × 1.2 NA water immersion objective was used for FCS. A 633 nm He−Ne laser was used for excitation. The laser power was set to 1% of the total laser power, corresponding to 20 μW. The emission detection was done with GaAsP spectral detector in the range of 650–700 nm.

Due to movement of cell membrane, the so-called “scanning FCS”^50^ modality was chosen to acquire data. Briefly, the line scan of 20 – 40 pixels was performed 50 000 times across the plasma membrane in the equatorial plane of the cell. Furthermore, to account for the cell movement, from every line scan the point with highest fluorescence intensity was chosen to represent this time point. The resulting trace was further correlated and fitted.

FCS fitting and diffusion coefficient calculation was performed using the home-made python based program Py-Profiler^51^. Curves were fitted with the following two-dimensional diffusion model:

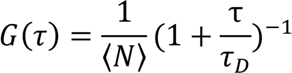

Where *G*(*τ*) is a correlation function, *τ* stands for delay time, *τ***_D_** – for diffusion time, *AR* – for aspect ratio of the focal volume and 〈*N*〉 - for an average number of molecules within the focal volume.

Calibration was performed using 10 nM solution of Alexa467(ThermoFisher) in pure water. Diffusion was calculated from the diffusion time using the following expression:

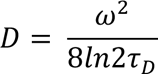

Where *D* is a diffusion coefficient, *ω*^2^ stands for full width at half maximum of focus point spread function, *τ*_*D*_ – for diffusion time. The trace extraction, correlation and fitting were performed using home-made open-source software called “Py_profiler”. The release with the sFCS functionality is available here: https://github.com/taras-sych/Single-particle-profiler/tree/sFCS_Release_v1.0

### Inner Leaflet analysis with NileRed fluorescence lifetime

2×10^6^Jurkat T cells (WT and KO) were transfected with SH4-HaloTag or GG-HaloTag. SH4-Halo is the N-terminal sequence of Lyn that codes for the myristol- and palmitoyl-acylation sites with a C-terminal HaloTag. GG-HaloTag is the C-terminal sequence of K-ras, which codes for the fatty acylation site and polybasic region. An N-terminal HaloTag was appended to this sequence. Both probes localize to the inner leaflet of the plasma membrane. 24 h after transfection, the cells were washed two times with Tyrode’s Buffer (25 mM HEPES, 150 mM NaCl, 5 mM KCl, 5.4 mM glucose, 1 mM CaCl2, 0.4 mM MgCl2, pH 7.2) and incubated with 200nM NR12-Halo (kindly provided by Andrey Klymchenko, University of Strasbourg) in Tyrode’s Buffer for 30 min at 37 °C. The cells were washed two more times in complete medium for 10 min at 37 °C and imaged immediately using a Leica SP8 confocal microscope with a 63x water immersion objective. FLIM images were obtained with 485 nm excitation and emission from 550–800 nm and fit using Leica LAS n-exponential reconvolution with two lifetimes. We report the intensity-weighted mean lifetime value.

### Proxiome analysis

Surface proteome mapping was performed using the molecular pixelation assay (MPX) by Pixelgen Techologies^32^. The Jurkat T WT cells or TMEM30A KO cells processing and subsequent NGS sequencing of the DNA libraries were performed according to the instructions provided in the MPX v2 User Manual (v2.00) by Pixelgen Technologies. Subsequent data analysis including the pixelator pipeline, quality control, abundance normalization, data integration, cell annotation as well as differential abundance, polarity and colocalization analysis was performed according to the tutorials by Pixelgen Technologies. Briefly, Cells were processed using the Pixelgen Single Cell Spatial Proteomics Kit (Pixelgen Technologies, PXGIMM001) following the manufacturer’s protocol and recommended reagents. Briefly, cells were thawed, washed, counted, and fixed with 1% PFA (Electron Microscopy Sciences, 15710). After blocking, each sample was divided into duplicates and stained with an 80-plex oligo-conjugated antibody panel, then stabilized using a secondary antibody. From each replicate, 20,000 stained cells were subjected to Molecular Pixelation. Molecular Pixels A were added first, followed by a gap-fill reaction and subsequent removal of Pixel A. Molecular Pixels B were then added and a second gap-fill reaction was performed. The cells were counted, and 1,000 cells from each replicate were processed with exonuclease treatment and indexing PCR. The resulting PCR products were pooled and purified using AMPure XP beads (Beckman-Coulter, A63881). Product purity was confirmed by gel electrophoresis on a TBE gel. The purified libraries were spiked with 15% PhiX (Illumina, FC-110-3001) and sequenced using paired-end sequencing (read 1: 28 cycles; read 2: 66 cycles; i7 index: 8 cycles; i5 index: 8 cycles) on an Illumina NextSeq 2000 system (donors 1 and 2: P3 flow cell; donor 3: P2 flow cell).

### Metalloprotease inhibitor treatment of Jurkat WT and TMEM30-KO

For metalloprotease inhibitor experiments, WT and TMEM30-KO Jurkat at 4 x 10^5^ cells/well of a U-bottomed 96 well plate in 200 µL RPMI/10% FBS were treated with either GI or GW at 10 µM overnight at 37 °C/5% CO_2_. Cells were centrifuged for 5 min at 400 x *g* at room temperature and resuspended in FACS buffer for subsequent antibody labeling.

### Flow cytometry for mucin analysis

WT and TMEM30-KO Jurkat were plated at 4 x 10^5^ cells/well of a U-bottomed 96 well plate in 50 µL 1 x annexin-V binding buffer (BD Biosciences 556454) for 20 min at room temperature, containing the following antibodies: anti-human CD43-PE clone 10G7 (Biolegend 343204) at 4 µg/mL; anti-human CD45-PE (MEM-28, Abcam ab134202) used at 1:100; anti-human MUC1-PE clone 16 A (Biolegend 355604) used at 4 µg/mL; anti-human MUC24-PE clone 67D2 (Biolegend 324808) used at 0.5 µg/mL; anti-human PSGL-1-PE clone TC2 (Invitrogen A15789) used at 1:100, CD62L-PE clone DREG-26 (BD Biosciences 555544) used at 1 µg/mL; annexin-V-FITC (Biolegend 640906) used at 0.9 µg/mL. All antibody labeling was done in conjunction with the corresponding concentration-matched isotype control antibody. After labeling, cells were washed 2 x with annexin-V binding buffer, centrifuged for 5 min at 400 x *g*, and resuspended in 2 % paraformaldehyde (Sigma-Aldrich 158127) for 15 min. After washing, cells were resuspended in annexin-V binding buffer and analyzed by flow cytometry using a Symphony A3 flow cytometer (BD Biosciences) and data processed using the FlowJo-V10 software (FlowJo, LLC). Gating on the relevant cell population was set according to Forward Scatter (FSC) and Side Scatter (SSC) before doublet exclusion.

## Data Availability

All data will be available in Karolinska Data Center upon publication.

## AI Use

No AI tools were used in this manuscript.

## Author Contributions

All authors performed the experiments. ES and CG performed microscopy and flow cytometry analyses. FR performed molecular pixelation experiments. LK and FT created the KO cell lines. MM performed flow cytometry for mucin expression analysis. IL and KRL performed leaflet specific biophysical experiments. ES, QS, IL, KRL and FT supervised the team and provided funding. All authors contributed to manuscript writing and editing.

## Acknowledgements

ES has been supported by Swedish Research Council Grants (grant no. 2020-02682, 2024-02993 and 2024-00289), Wellcome Leap’s Dynamic Resilience Program (jointly funded by Temasek Trust), Cancer Fonden (25 4592 Pj) Karolinska Institutet (2024-03250; 2024-03341; 2022-00803; 2020-00997), Cancer Research KI (2024-03488), Strategic Research Programme in Diabetes at Karolinska Institutet, Human Frontier Science Program (RGP0025/2022), Longevity Impetus Grant from Norn Group, Hevolution Foundation and Rosenkranz Foundation. ES is an EMBO Young Investigator (YIP-2025). COG and FR were supported by Karolinska Institutet Doctoral funding (KID, 2022-00803, 2020-00997). We thank the SciLifeLab Advanced Light Microscopy facility and National Microscopy Infrastructure (VR-RFI 2016-00968) for their support on imaging and Pixelgen on their support for the molecular pixelation assay.

