## Supplementary Figures for "Cell surface remodeling caused by the loss of flippase subunit TMEM30A in immune cells"

**for**

### Supplementary Figures:

A

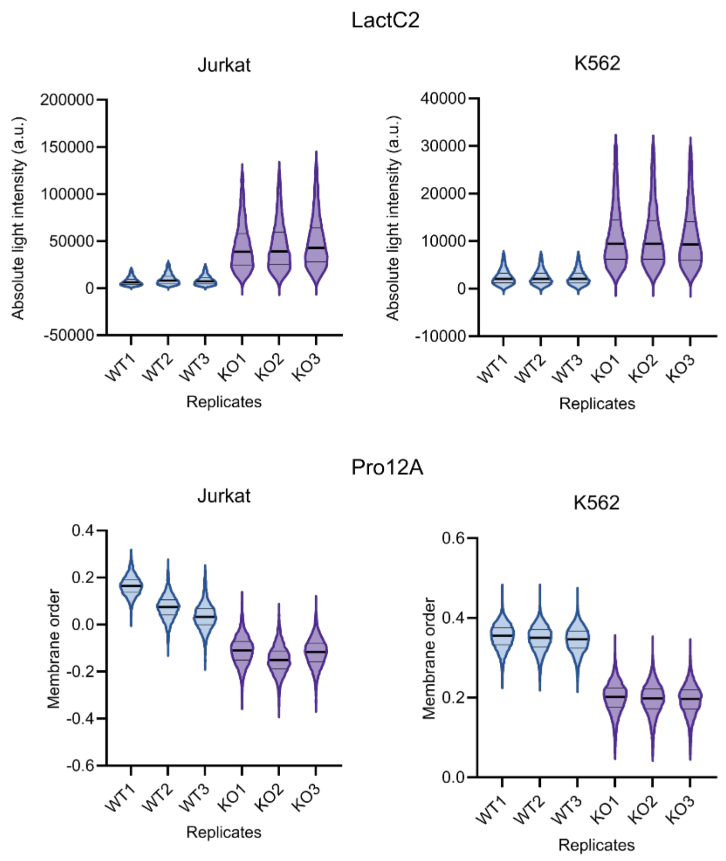


**Supplement Figure, SF1.** A) Absolute light intensity measurements of LactC2 and B) membrane order of Pro12A for all replicates with spectral flow cytometry.

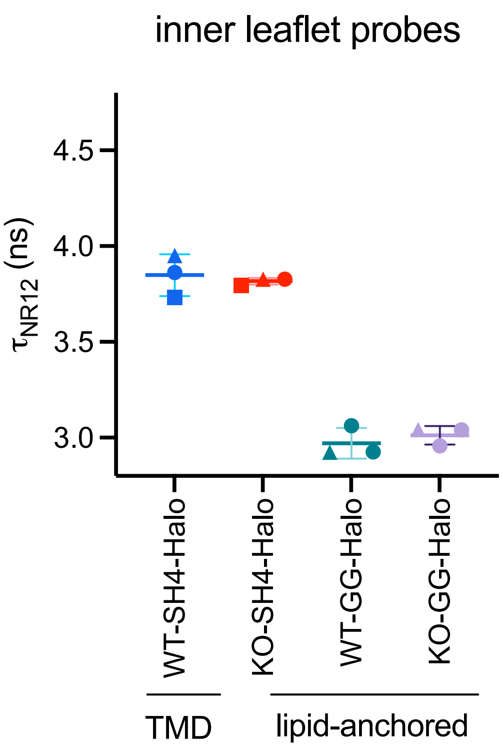


**Supplement Figure, SF2.** Lipid order in the inner leaflet does not change notably. HaloTag-Nile Red was used to measure local lipid environment of three reporter proteins: one ordered domain (SH4) and one disordered domain (GG) preferring. Lifetime of Nile Red is shown for three peptides. Each point is a replicate.

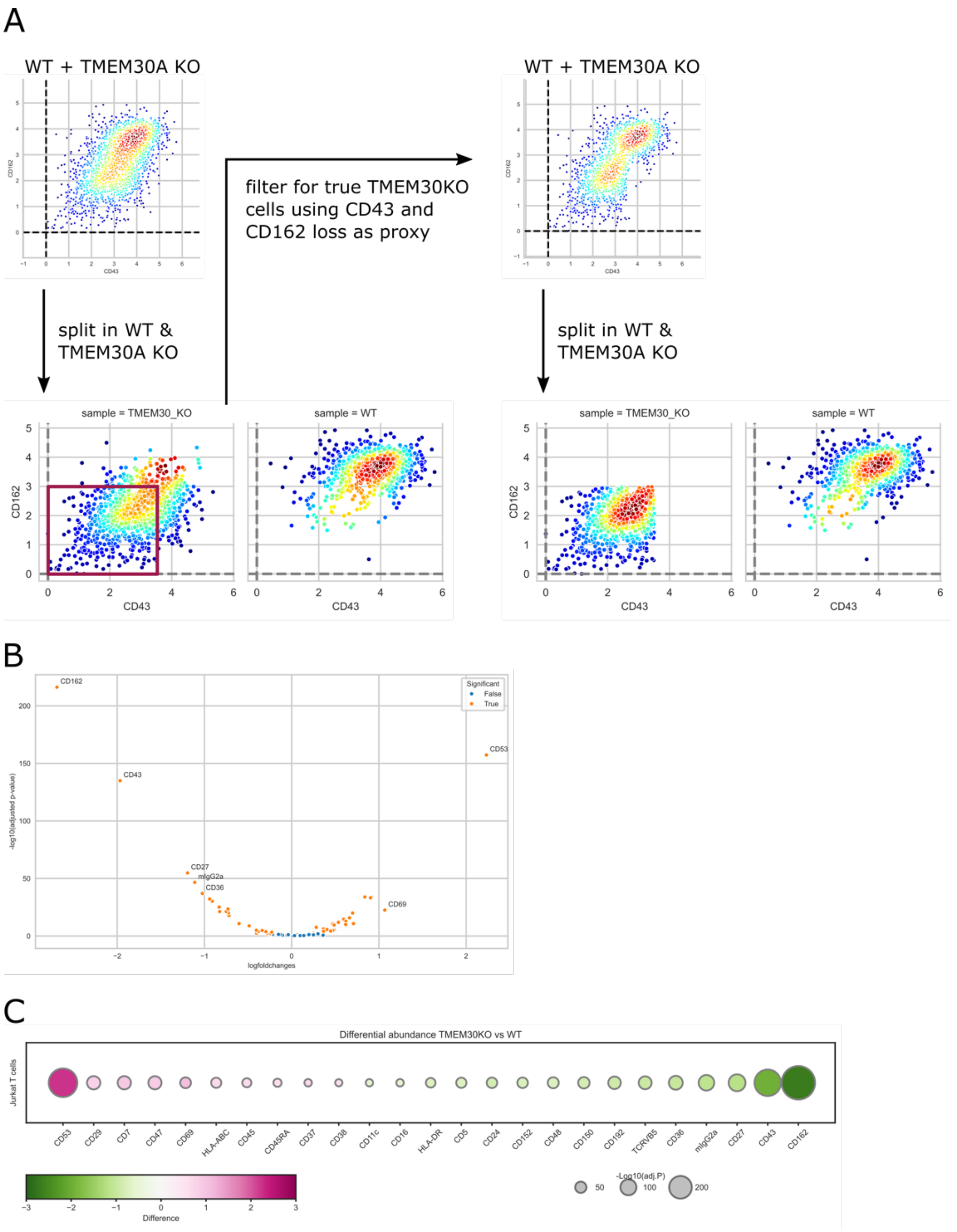


**Supplement Figure, SF3:** Surface protein abundance after filtering for true TMEM30A KO cells. A) TMEM30A KO cells are a cell pool and not all cells have the KO as indicated by cells expressing CD43 and CD162. TMEM30A KO cells are filtered for cells with a shifted clr value of CD162<3 and CD43>3.5. After filtering two distinct cell populations are visible. B) Volcano plot for the abundance of proteins differentially regulated in WT vs. true KO Jurkat cells. Orange dots indicate significantly up- or downregulated proteins (adjusted p-value < 0.01). Labels are added for proteins with a log fold change > 1. C) Proteins of differential abundance (log fold change >0.6) in WT vs. true KO Jurkat cells. Color corresponds log fold change and size of the bubbles indicates the adjusted p-value.

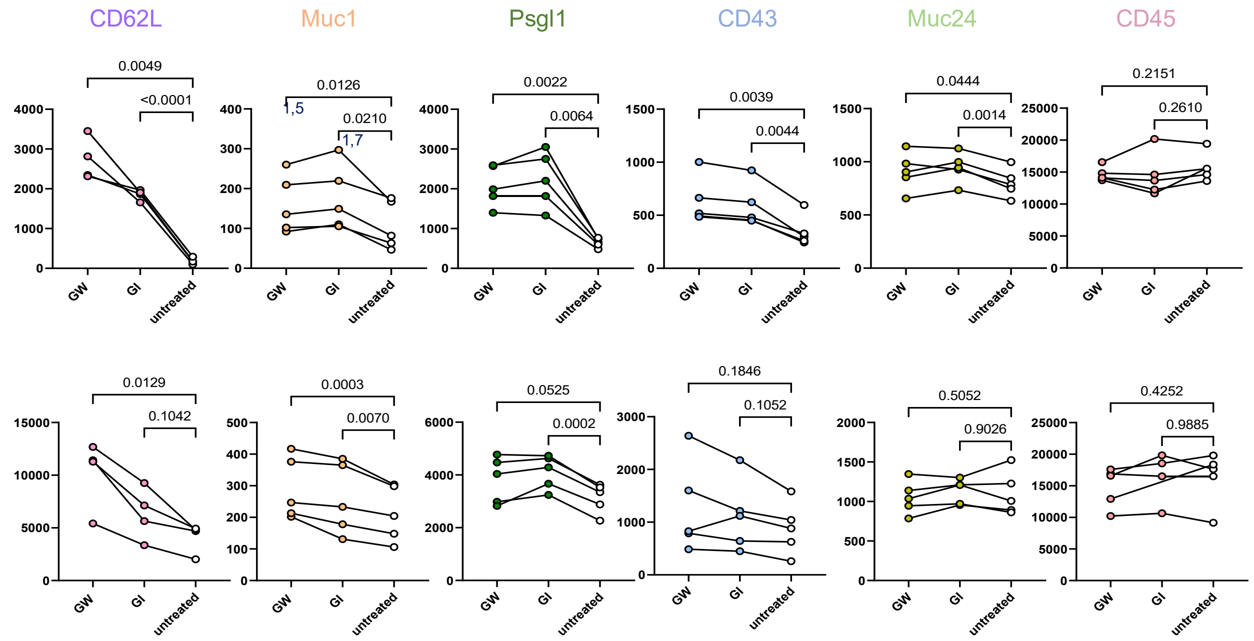


**Supplementary Figure, SF4.** The protein shedding by ADAM10 in Jurkat WT and TMEM30A-KO cells. Top panel shows the TMEM30A-KO cells and bottom panel is WT cells. ADAM10 inhibitor GI254023X (GI) and ADAM family inhibitor GW280264X (GW) reduces the shedding significantly in TMEM30A-KO cells and notably in WT cells.
